# Expression and purification of human neutrophil proteinase 3 from insect cells and characterization of ligand binding

**DOI:** 10.1101/2023.11.10.566610

**Authors:** Fahimeh Khorsand, Bengt Erik Haug, Inari Kursula, Nathalie Reuter, Ruth Brenk

## Abstract

Neutrophil proteinase 3 (PR3) is an important drug target for inflammatory lung diseases such as chronic obstructive pulmonary disease and cystic fibrosis. Drug discovery efforts targeting PR3 require active enzyme for *in vitro* characterization, such as inhibitor screening, enzymatic assays, and structural studies. Recombinant expression of active PR3 overcomes the need for enzyme supplies from human blood and in addition allows studies on the influence of mutations on enzyme activity and ligand binding. Here, we report the expression of recombinant PR3 (rPR3) using a baculovirus expression system. The purification and activation process described resulted in highly pure and active PR3. The activity of rPR3 in the presence of commercially available inhibitors was compared with human PR3 by using a fluorescence-based enzymatic assay. Purified rPR3 had comparable activity to the native human enzyme, thus being a suitable alternative for enzymatic studies *in vitro*. Further, we established a surface plasmon resonance-based assay to determine binding affinities and kinetics of PR3 ligands. These methods provide valuable tools for early drug discovery aiming towards treatment of lung inflammation.

## Introduction

Lung inflammation which occurs in diseases such as asthma, chronic obstructive pulmonary disease (COPD), cystic fibrosis (CF), and pneumonia is a serious challenge to public health worldwide (1). COPD, mainly caused by environmental factors, such as smoking, is projected to be the third leading cause of death by 2030 (2,3). Airway epithelial dysfunction and airway neutrophilic inflammation leading to airflow obstruction and impaired lung function are involved in the initiation and progress of clinical conditions associated with COPD and CF (4). As neutrophils migrate in response to an infection, serine proteases are released from their azurophil granules and play a major role in degrading proteins from pathogens but also target the extracellular matrix proteins to facilitate neutrophil migration (5). Imbalance between these serine proteases and their endogenous inhibitors and increased protease activity have a prominent role in the destruction of lung parenchyma (emphysema) (6) and as a result, neutrophil serine proteases have been in focus as potential drug targets for the treatment of inflammatory lung diseases.

The roles of the medically important serine proteases neutrophil elastase (hNE), cathepsin G (CatG), and proteinase 3 (PR3) have been characterized in neutrophils. They are involved in a variety of inflammatory human conditions, including chronic lung diseases, in addition to their involvement in pathogen destruction and the regulation of proinflammatory processes (7). hNE has been the subject of drug discovery studies for over 30 years (see e.g.(8,9)). Several inhibitors have been developed, and some of them have been progressed to clinical studies, but only sivelestat (ONO-5046) has been approved in some countries (Japan and South Korea) for the treatment of acute lung injury and respiratory distress (10,11). Studies on some hNE inhibitors, such as ONO-6818, alvelestat (AZD9668), and AZD6553, have been discontinued owing to unsatisfactory results from clinical trials (12). BAY-85-8501 is a new highly potent hNE inhibitor and has been evaluated in a phase II trial for safety and efficacy in patients with non-cystic fibrosis bronchodilation (13,14). While studies on hNE are still in progress, another neutrophil serine protease, proteinase 3 (PR3), has drawn the attention as a drug target over the recent years. Despite PR3 and hNE being homologues and sharing 56% sequence identity, their ligand binding sites exhibit notable differences. The two enzymes have therefore slightly different ligand and inhibitor selectivity, and existing drug candidates targeting hNE are not necessarily also potent inhibitors of PR3 resulting in selective inhibition of some compounds (7,15). Yet, PR3 is a promising drug target for COPD and CF for several reasons: higher levels of PR3 than hNE are released from neutrophil azurophil granules, PR3 is active as both a membrane-bound and a free enzyme, PR3 is present in a larger area of the cell than hNE and local lung derived inhibitors have less impact on PR3 than hNE (16–18).

Owing to the importance of evaluating the potency of drug candidates against PR3, preparing a sufficient amount of this enzyme for crystallization, binding assays, and activity screening is in high demand. PR3 purification from neutrophils is a difficult and costly process with low yields. Expression of recombinant PR3 (rPR3) in bacteria (19), yeast (19), insect cells (20,21), human mast cells (22), and adherent 293 cells (23) have been used to study the recognition of rPR3 by anti-PR3 autoantibodies (PR3-ANCA) or to study the processing of the amino- and carboxy-termini. However, production of rPR3 for activity assays has not been reported.

Here, we aimed at to expressing active PR3 suitable for kinetic experiments and ligand studies. We report the expression of rPR3 using *Spodoptera frugiperda* (Sf9) and the results of inhibition assays for rPR3 and PR3 purified from neutrophils to compare the activities of these proteins. As information on binding kinetics is of important for drug discovery (24), we also developed a surface plasmon resonance (SPR) assay for the determination of binding affinities and kinetics of rPR3 inhibitors. The developed methods provide an experimental framework to support drug discovery efforts for PR3.

## Methods

### Chemicals and reagents

All chemicals and reagents were of analytical grade. The elastase inhibitors selected for the inhibition study of rPR3 were purchased from the following suppliers: α1-antitrypsin (Sigma-Aldrich), sivelestat (Sigma-Aldrich), alvelestat (MedChemExpress), and BAY-85-8501 (MedChemExpress). Enzymes purified from human neutrophiles (hPR3 and hNE) were purchased from Athens Research and Technology. hNE substrate (MeOSuc-AAPV-AMC) was supplied by Santa Cruz Biotechnology. The FRET peptide used as the PR3 substrate in enzymatic assay was prepared as described before (25).

### Plasmids/vector constructs

Five constructs (P2-6, Figure 1) were synthesised and cloned in to the pFastBAC1 vector by Genscript. The Mellitin signal sequence to direct secretion of the expressed protein (26) was inserted upstream of the PR3 sequence (either the sequence for the whole protein or the mature protein) followed by a TEV cleavage site and either a His-tag or Strep-tag (see supplementary material for the DNA sequences of the different constructs).

**Fig 1.**
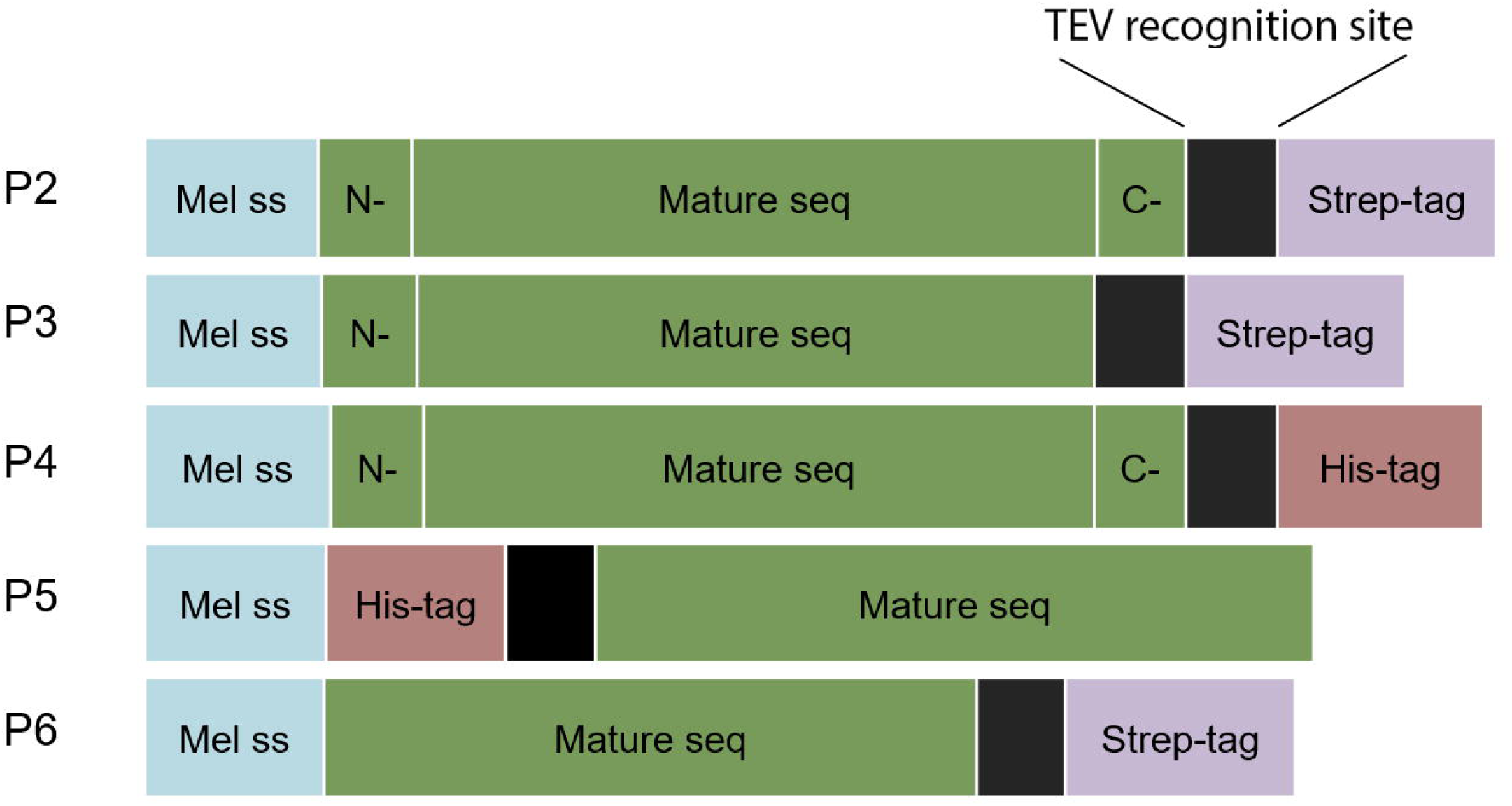
The different rPR3 constructs used in this study. A melittin signal sequence (*Mel ss,* marked as blue box) and a TEV cleavage site (black box) were included in all constructs. Three constructs contained a C-terminal Strep-tag (P2, P3, and P6, purple box), a C-terminal His-tag (P4, pink box) or an N-terminal His-tag (P5). In addition to the sequence for the mature PR3 (green box), three constructs contained the N-terminal pre-peptide (P2-4, N-in green box) and two the C-terminal propeptide (P2 and P4, C-in green box).

### Bacmid and baculovirus preparation

Bacmids were generated using the EMBacY MultiBac system (Geneva Biotech). Genes of interest were inserted into bacmids by Tn7 transposition. DH10EMBacY competent *E. coli* (EMBL, Grenoble), which contained both the EMBacY genome (bacmid) and the helper plasmid (IPTG inducible Tn7 transposon complex with tetracycline resistance gene), were transformed with pFastBac1 vectors containing the genes for the different PR3 constructs and plated onto selection agar plates. Positive colonies were selected based on blue/white screening and the bacmid was purified for transfection and virus production in Sf9 insect cells. Bacmids containing the PR3 sequences were isolated and checked by PCR with M13 primers to make sure that the size of the bacmids was the same as expected (see supplementary material, Fig S1a).

### Expression of rPR3

Recombinant protein was expressed in baculovirus-infected insect cells using the protocol described by Bieniossek *et al.* (27). Sf9 cells (Invitrogen) were transfected with the bacmid, using the FuGENE®6 transfection reagent (Promega), and virus particles were harvested after 7 d of infection. The recombinant primary virus was amplified to a high-titre viral stock. Approximately 9 µl (1 × 10^6^ cells) of the high-titre virus were used to infect the insect-cell culture at a cell density of 0.6–0.8 × 10^6^ cells/mL. The culture was centrifuged at 7000 g for 30 min after 5 d, and the supernatant (clarified expression medium) was filtered with 0.45 µm sterile filters and used for protein purification.

### Monitoring of rPR3 expression

To monitor expression of rPR3, 15 mL samples were collected 24, 48, and 72 h after the Sf9 cells infected with baculoviruses made with the different constructs reached growth arrest. The culture was centrifuged at 1000 g for 15 min, the cleared medium was concentrated 30 times to the final volume of 500 μL and was stored at −20 °C. The pellet was resuspended in 300 µL PBS and stored at −20 °C. To perform the dot blot, samples were thawed. The cells were centrifuged to remove PBS and resuspend with the lysis buffer (50 mM Tris-HCl, pH 8 with 50 mM NaCl, protease inhibitor). 2 µL from the supernatant and cell lysates were applied on a nylon membrane and two different antibodies (an HRP-conjugated anti-His tag from Qiagen and an HRP-conjugated anti-strep tag from Sigma-Aldrich) were used on separate blots to detect PR3. The metal enhanced DAB substrate kit (ThermoScientific) was used for staining of the blots.

### Purification

The supernatant from 5-10 L culture was loaded on an HisTrap Excel column (Cytiva) overnight and washed for 20-30 column volumes with buffer A (50 mM potassium phosphate, 150 mM NaCl, pH 7.4). Another washing step was carried out with buffer B (buffer A plus 50 mM imidazole) for 10 column volumes, and the protein was eluted with a gradient of 50 to 400 mM imidazole in buffer A over 10 column volumes. The eluted protein was concentrated and loaded on a SEC200 increase 10/300 column (Cytiva), and was eluted using buffer C (50 mM sodium acetate, 150 mM NaCl, pH 5.5). The identity of the purified protein was confirmed using mass spectrometry (Fig S1c).

### Activation of rPR3

To cut off the N-terminal dipeptide from PR3 and activate the enzyme, purified rPR3 and cathepsin C (Sigma-Aldrich C8511) were mixed at a ratio of 95:5 in the activation buffer (5 mM mercaptoethanol in buffer C) and incubated at 35 °C for 18 h.

### Enzymatic activity assay

The enzymatic assay is similar to the one we reported earlier (28) with minor modifications. Briefly, to determine the inhibition of PR3, 0.25 nM of enzyme was incubated with buffer (50 mM HEPES pH 7.4, 750 mM NaCl, 0.5% Igepal and 1% DMSO) or buffer containing the inhibitors at various concentrations for 30 min at RT. The reaction was started by adding the FRET peptide (Abz-VADnVADYQ-EDDnp) at a final concentration of 5 μM. The fluorescent signal (excitation filter 320, emission filter 420) was read every 30 s for 30 min at 30 °C with a fluorescent plate reader (Tecan Spark) and the enzymatic activity was measured from the initial linear portion of the slope (fluorescent signal/min) of the time-course curve of reaction progress.

For the activity assay with hNE, 0.25 nM of enzyme was incubated as described above for PR3. The reaction was started by adding the fluorogenic substrate (MeOSuc-AAPV-AMC) at a final concentration of 5 μM. The fluorescent signal (excitation filter 360, emission filter 460) was read as described above for PR3.

Data were fitted using nonlinear regression analysis (GraphPad Prism v.9) to determine the IC_50_ values.

### Biotinylation of rPR3

50 μL activated rPR3 (2 mg/mL) in PBS was mixed with NHS-PEG4-Biotin at a molar excess of 20-fold and incubated at RT for 30 min. Unreacted biotin was removed by 4 times diluting and concentrating the buffer so that the final concentration was 1/10000 of the starting concentration. Biotinylation was confirmed using a chromogenic detection kit (Thermo ScientificTM).

### SPR assay

The SPR data were obtained using a Biacore T200 (GE Healthcare Life Sciences) and presented as the mean of triplicate measurements. Briefly, 100 μL of 200 nM biotinylated rPR3 in PBS was injected and captured on a NAHLC 1500M chip (XanTec Bioanalytics GmbH) at a flow rate of 5 μL/min. Biotinylated protein was captured to an immobilization level of around 4000-8000 RU. As reference, an unrelated protein available in our group, biotinylated δMtDXR (*Mycobacterium tuberculosis* 1-deoxy-D-xylulose 5-phosphate reductoisomerase) expressed in BL21(DE3) *E. coli* cells and biotinylated enzymatically with GST-BirA, was immobilized on the reference flow cell (3500-4000 RU). The running buffer for the immobilization of the proteins was PBSP (GE Healthcare Life Sciences). The binding assay was carried out using the same buffer and a flow rate of 30 μL/min at 25 °C. To measure compound binding, different concentrations of the studied compounds were injected for 200-300 s over the flow cells starting with the lowest concentration and dissociation was initiated with the injection of the running buffer which continued for 600-900 s. For regeneration, 1M NaCl was injected for 1 min. Sensorgrams were double-referenced by subtracting the signal from a reference surface and the signal from one blank injection. The Biacore T200 Evaluation Software 3.0 was applied for the determination of steady state affinity and all the kinetic data using a global fit. K_D_ values were determined using an equation corresponding to a reversible, one-step, 1:1 interaction model and fitting values taken at the end of the injection (representing steady-state signals) by nonlinear regression analysis.

## Results and discussion

### rPR3 expression and purification

As a baculovirus expression system has been previously used to produce recombinant PR3 which yielded the only available crystal structure so far (20), and as a eukaryotic expression system is needed for posttranslational modifications (mainly glycosylations) and for the removal of preprosequences, we opted for using *Spodoptera frugiperda* (Sf9) for recombinant expression of PR3. Five different constructs were made for the expression of rPR3 (Fig 1, see Supporting information for sequences). The mature PR3 sequence lacking both the N- and C-terminal preprosequences and the Ala-Glu dipeptide from the propeptide was used in the constructs P5 and P6 while the whole PR3 sequence was inserted in the constructs P2-4. The N-terminal propresequence of PR3 functions as the endogenous signal peptide. All constructs contained a melittin signal peptide at the N-terminus to direct the protein through the post-translational modification and secretory pathway as previously described (19). Two types of affinity tags were investigated, a 6x His-tag either at the N-terminal (P5) or at the C-terminal (P4 and P7), and a C-terminal Twin-Strep-tag (P2, P3 and P6). All constructs also contained a TEV cleavage site between the PR3 sequence and the affinity tag to enable cleavage of the tag if required.

Expression of the different constructs was monitored by measuring the fluorescence of YFP (yellow fluorescent protein) which was co-expressed in the insect cells. YFP became visible after around 48 h post-infection indicating that infection had been successful for all constructs (Fig S1b).

To determine if the different proteins were secreted into the media, we used anti-His and with anti-Strep-tag antibodies. For the constructs P2, P3, P4 and P6 protein was clearly detectable in the lysate and for P2, P3 and P4 protein was also detectable in the media (Fig 2). This indicates that the N-terminal preprosequence of PR3 (endogenous signal peptide) is essential for the secretion of protein to the medium. As the post-translational modifications are associated with the secretory pathway, we considered secreted rPR3 to be of higher interest than non-secreted rPR3. Accordingly, the constructs lacking the N-terminal preprosequence of PR3 (P5 and P6) were not considered for further studies. Given the high cost associated with using Strep-Tactin columns for purification of Strep-tag containing proteins and apparently similar expression levels for secreted proteins containing either a Strep-tag (P2 and P3) or a His-tag (P4), we decided to continue further work with the construct containing a His-tag.

**Fig 2.**
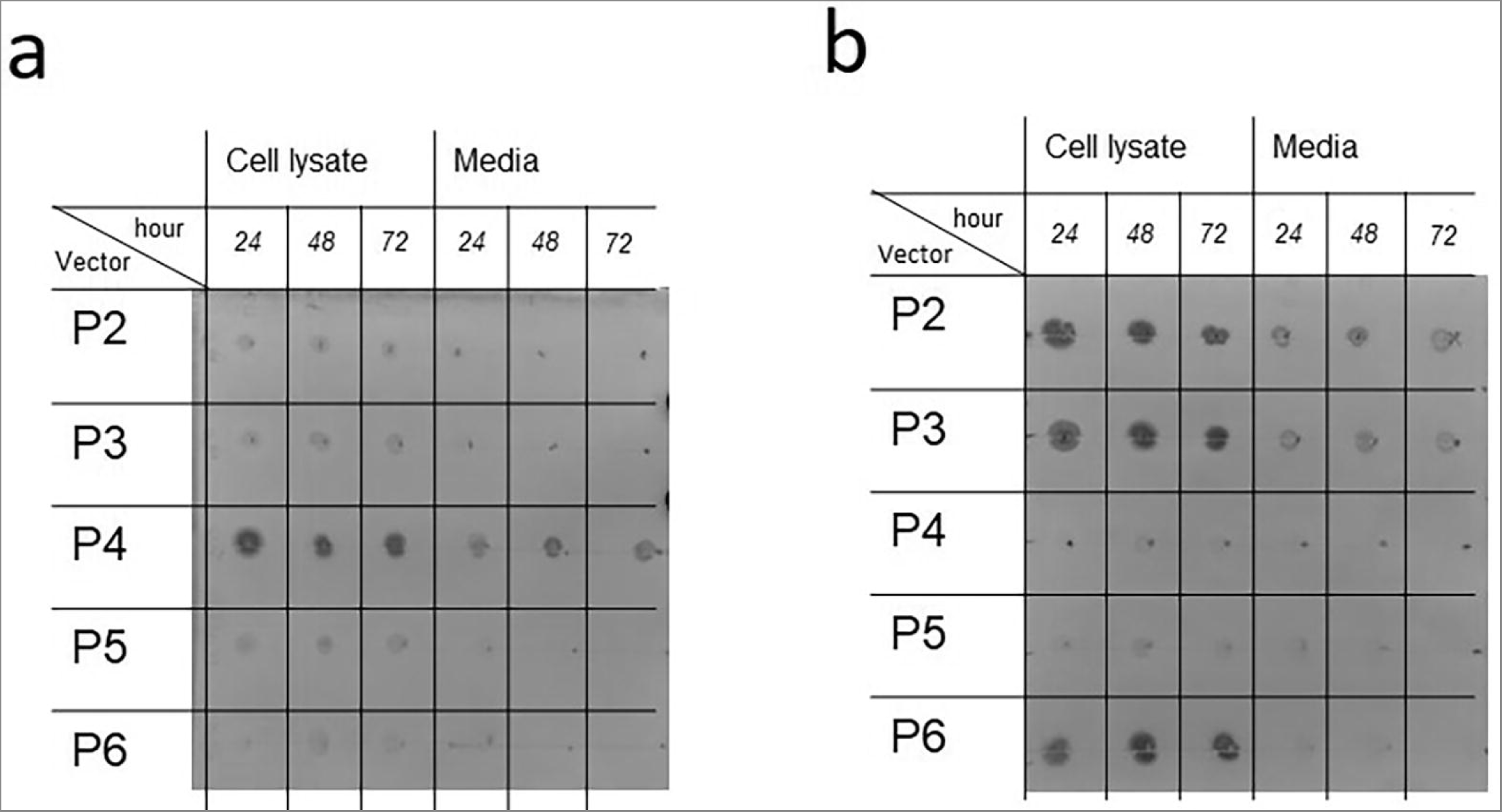
Dot blot analysis of rPR3 expression. The time given refers to the time the sample was taken after cell growth arrest was reached. Blot a was treated with an anti-His antibody and blot b with and anti-Strep-tag antibody.

Subsequently, rPR3 obtained using construct P4 was purified. Purification of rPR3 with a His-trap column resulted in a purity of approximately 80% (Fig. 3). Therefore, size exclusion chromatography column was subsequently used to obtain pure rPR3. The purity of rPR3 was very high at this stage (Fig. 3) and the protein yield was around 400 μg/L of culture medium. The identity of rPR3 was confirmed using mass spectrometry (Fig S1c and d).

**Fig 3.**
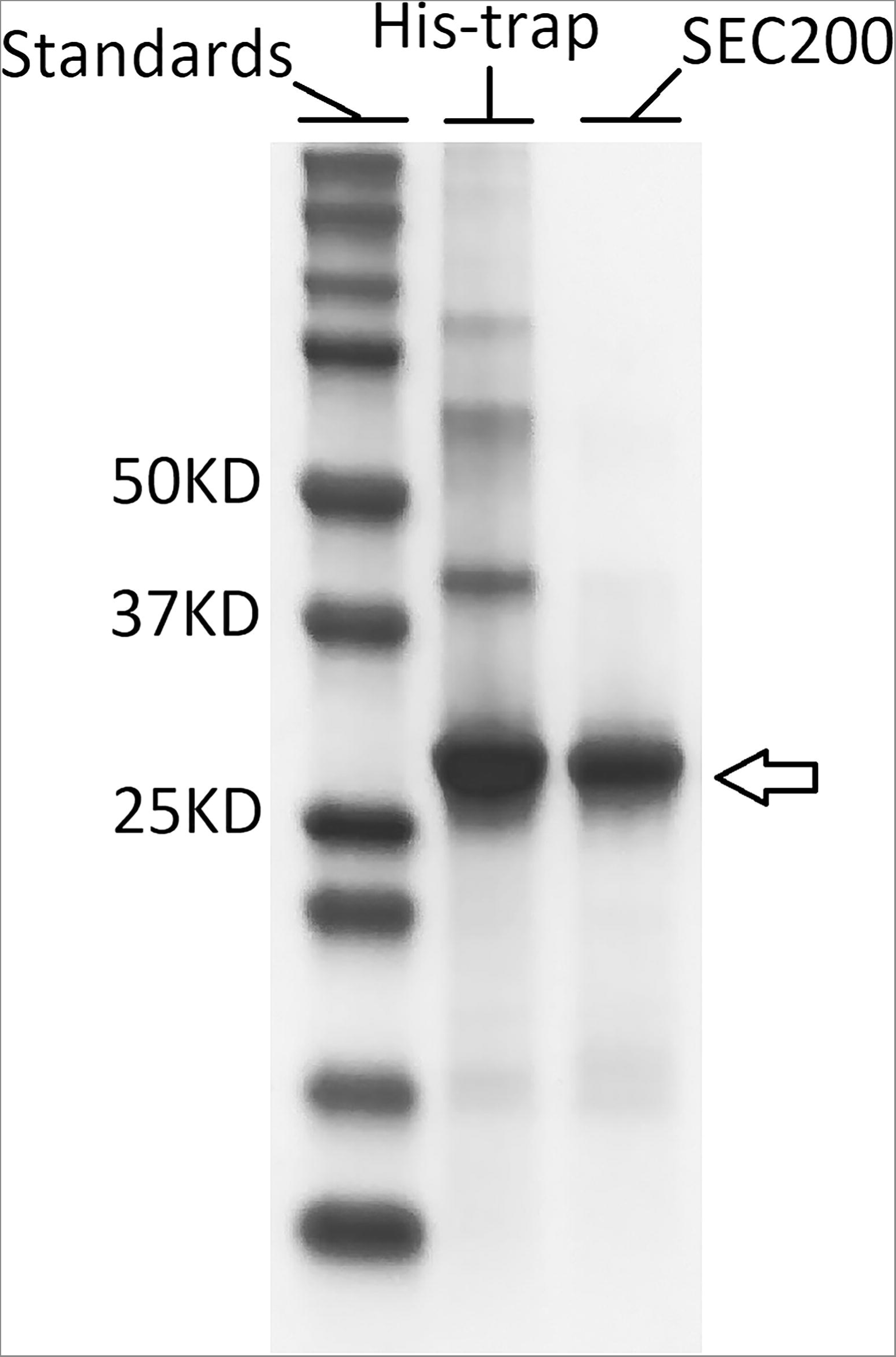
SDS-PAGE showing the purity of rPR3. Purification was carried out with an Excel His-trap column (lane His-trap) and in addition with a SEC200 column (lane SEC200).

In the next step, rPR3 was activated and the activity of the mature enzyme was assessed. To activate rPR3, the N-terminal dipeptide needs to be removed (20). Here, this was achieved by treating PR3 with cathepsin C. To determine the activity of the mature enzyme, a kinetic assay was conducted. Due to ease of use, we opted to use an activity assay with a fluorogenic substrate. However, it is not possible to determine kinetic parameters with this assay as this would lead to a too high background fluorescence signal. Therefore, we have assessed the activity by comparing IC_50_ values of some potent elastase inhibitors determined with hPR3 and rPR3 (Table 1). One of these inhibitors is α1-antitrypsin, an endogenous serine protease inhibitor, also denoted SERPIN, which plays an important role in balancing the activity of serine proteases in the body (29). The other three inhibitors are small molecule hNE inhibitors that have been developed by various pharmaceutical companies over recent years. For some of the compounds no inhibition data for PR3 is available. To allow comparison to literature values, we therefore determined also the IC_50_ values of these compounds for hNE (Table 1). Indeed, the obtained values of these inhibitors against hNE were comparable with the data available in literature giving confidence in the assay setup (Table 1, Fig S2). For alvelestat, enzyme inhibition could not be determined up to 2.5 μM. Measuring higher concentrations was not possible due to high background fluorescence. For the remaining compounds, very similar IC_50_ values for hPR3 and rPR3 were obtained (Table 1, Fig S2). These data show that we were able to produce rPR3 with similar activity as hPR3.

**Table 1.**
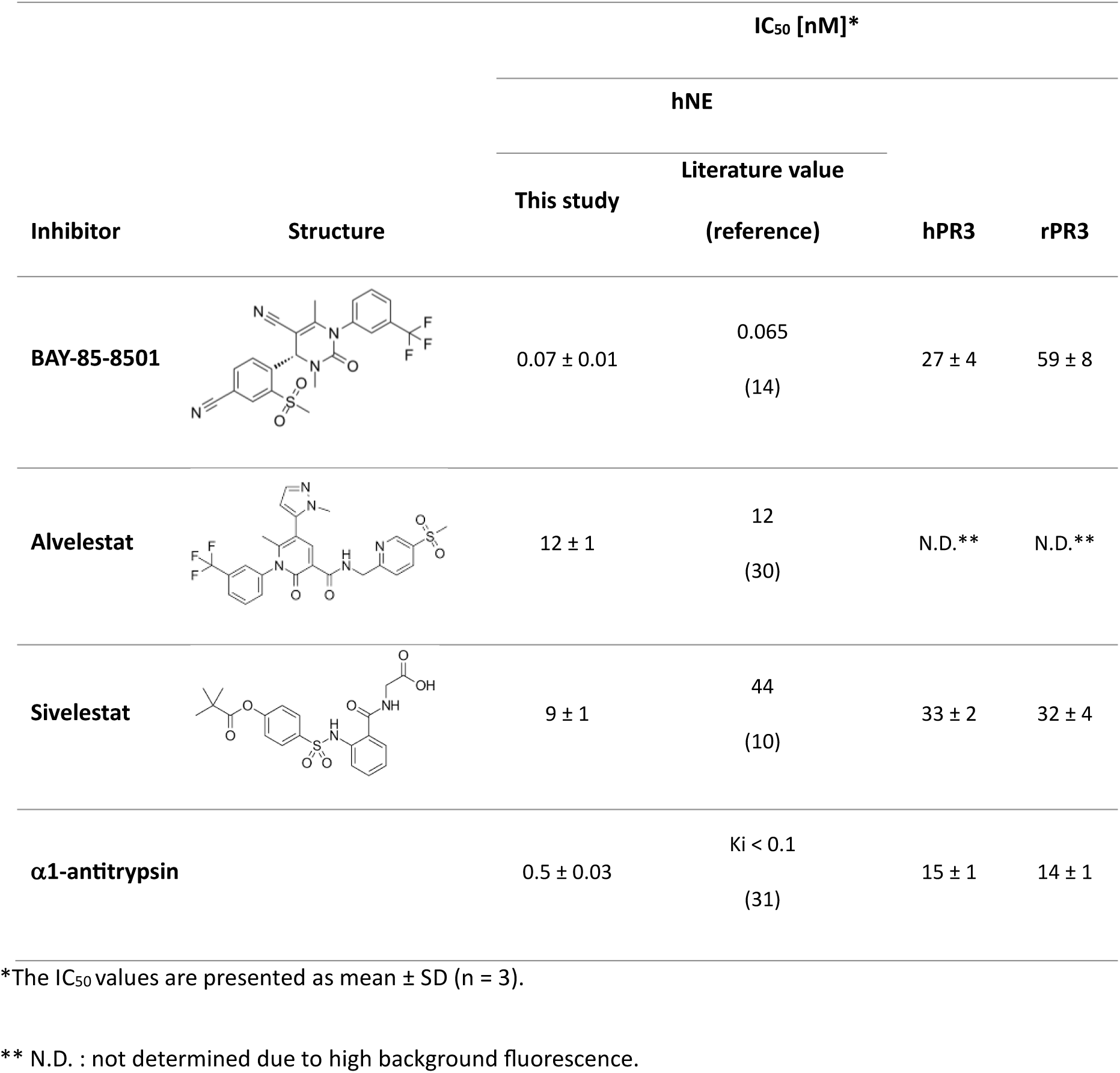
IC_50_ values for known elastase and PR3 inhibitors determined using hPR3, rPR3 and hNE.

| Inhibitor | Structure | IC<sub>50</sub> [nM]* |  |  |  |
| --- | --- | --- | --- | --- | --- |
|  |  | hNE |  |  |  |
|  |  | This study | Literature value | hPR3 | rPR3 |
|  |  |  | (reference) |  |  |
| **BAY-85-8501** |  | 0.07 ± 0.01 | 0.065<br>(14) | 27 ± 4 | 59 ± 8 |
| **Alvelestat** |  | 12 ± 1 | 12<br>(30) | N.D.** | N.D.** |
| **Sivelestat** |  | 9 ± 1 | 44<br>(10) | 33 ± 2 | 32 ± 4 |
| **α1-antitrypsin** |  | 0.5 ± 0.03 | Ki < 0.1<br>(31) | 15 ± 1 | 14 ± 1 |

\*The IC<sub>50</sub> values are presented as mean ± SD (n = 3).
\*\* N.D. : not determined due to high background fluorescence.

### SPR assay to determine binding affinities and kinetics of PR3 inhibitors

Next, we developed a binding assay for rPR3 using SPR. This method allows for determination of both binding affinities and kinetic parameters. Further, the SPR read-out is not influenced by fluorescence interference from test compounds, thus this method should therefore also allow for measurement of fluorescent compounds like alvelestat (Table 1). The assay was developed using neutravidin coated SPR chips. Neutravidin is a non-glycosylated analogue of avidin (32) and allows for capture and immobilization of biotinylated ligands onto the SPR chip. Accordingly, rPR3 was biotinylated with NHS-PEG4-biotin which reacts with primary amines such as the side chain of lysine residues or the amino-termini of polypeptides. The injection of biotinylated rPR3 into the flow cell led to high and stable binding to the chip (Fig S3). As reference, an unrelated protein available in our laboratory, biotinylated δMtDXR was selected and immobilized in the reference cell (Fig S3). The binding sites of rPR3 and DXR are very different, thus affinity of the investigated rPR3 inhibitors towards DXR was not expected. The inhibitors were injected in a range of concentrations and binding responses were analysed (Fig 4).

**Fig 4.**
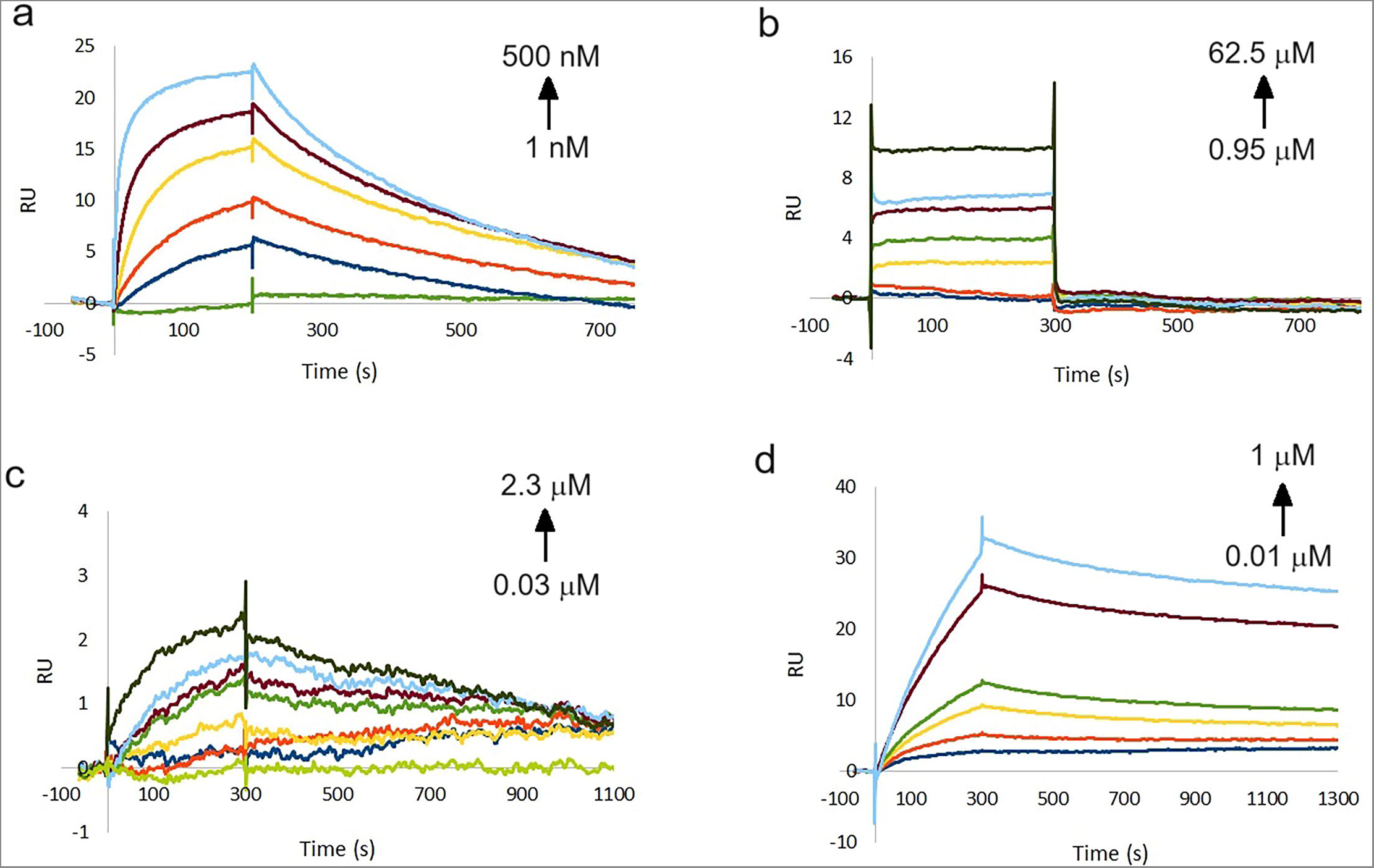
Sensorgrams of (a) BAY-85-8501, (b) alvelestat, (c) sivelestat, and (d) α1-antitrypsin binding to rPR3.

For BAY-85-8501, a K_D_ of 47 nM was determined using steady state kinetics and a K_D_ of 14 nM using flow kinetics (Fig 4a, Fig S3a, Table 2). These values are comparable to the K*_i_* value of 50 nM derived from the IC_50_ value for this compound (Table 1) using the Cheng-Prusoff equation (K_i_ = IC_50_/(1+ [S]/K_m_) and the literature K_M_ value of 27.4 µM (25). Using the 1:1 binding kinetic model, k_on_ and k_off_ rates were also obtained (Table 2). Using these values, the residence time of this molecule was calculated to be 5.2 min.

**Table 2.** Binding affinities and kinetic parameters of rPR3 inhibitors determined using SPR.

| Compound | $K_D$ ($\mu\text{M}$) | $k_{on}$ (1/Ms) | $k_{off}$ (1/s) | Steady state $K_D$ ($\mu\text{M}$) |
| --- | --- | --- | --- | --- |
| **BAY-85-8501** | $(0.14 \pm 0.03) \times 10^{-1}$ | $(2.25 \pm 0.65) \times 10^{+5}$ | $(3.19 \pm 1.5) \times 10^{-3}$ | $0.03 \pm 0.01$ |
| **Alvelestat** | N.D. | N.D. | N.D. | $13.5 \pm 5.6$ |

N.D. not determined because the data could not be fitted.
The reported values are the averages and SDs of triplicates.

When testing alvelestat in the SPR assay, we found it to be a weak rPR3 binder with fast association and dissociation rates (Fig 3b). Therefore, the binding constant could only be determined using the steady state method (Fig S3b) resulting in a K_D_ value of 13.5 μM (Table 2). This value is about 100 times higher than the IC_50_ value determined for the same compound with hNE while no affinity could be determined using the enzymatic assay (Table 1). The observed selectivity profile is in line with previous data (30).

We found that sivelestat and α1-antitrypsin hardly dissociated from rPR3 (Fig4c and d), which is consistent with the reported covalent binding capabilities of both these inhibitors (33,34). Inhibition of PR3 by α1-antitrypsin is reported to take place in at least two steps: the enzyme and the inhibitor first form a high-affinity reversible inhibitory complex EI* with an equilibrium dissociation constant K*i of 38 nM and then EI* subsequently transforms into an irreversible complex EI with a first-order rate constant k2 of 0.04 s^−1^ (34). Sivelestat has been reported to covalently inhibit hNE via acylation through formation of a pivaloyl ester (30).

## Conclusion

Here, we report a protocol for expression of rPR3. The protein was subsequently used for determination of ligand binding affinity and kinetics with SPR and inhibitor potency with a fluorescence-based inhibition assay. We have demonstrated that the activity of rPR3 is comparable with the wild-type human neutrophil PR3 making it suitable to study ligand binding and inhibition. Our protocol is also suitable to produce mutant variants of PR3 which will allow further investigation of PR3 structure-activity relationship in inhibitor binding. Such studies will be highly valued to shed light on what drives ligand affinity and selectivity, as for example here observed for alvelestat (Tables 1 and 2), especially as no crystal structures of PR3-inhibitor complexes are currently available. We have also developed an SPR-based protocol which can be used to determine binding affinity (K_D_) and kinetic constants The SPR data was in line with the inhibition data obtained using a fluorogenic enzymatic assay confirming the efficacy and accuracy of the SPR data to characterize ligand binding. Collectively, the reported methods will we very valuable tools to support drug discovery endeavours towards PR3.

## Acknowledgement

The work was supported by the Research Council of Norway (RESPOND3, grant number 294594). We made use of the Facility for Biophysics, Structural Biology and Screening at the University of Bergen (BiSS), which has received funding from the Research Council of Norway (RCN) through the NORCRYST (grant number 245828) and NOR-OPENSCREEN (grant number 245922) consortia. We would like to thank Bruna Schuck de Azevedo for providing us with the biotinylated MtDXR used in the SPR assay. MS data was collected at the Mass spectrometry core facility at the University of Oulu.

## Supporting information

All the DNA sequences and the corresponding protein sequences for the vector constructs P2-6 can be found in the supplementary material. The M13 primers sequence is also available in the supplementary material file. Additionally, the supporting information contains the four following figures:

**Fig S1 - Expression and identification of rPR3 in insect cells.** a) Agarose gel to detect bacmid production. b) Sf9 cells infected with baculovirus showing expression of YFP as green dots using fluorescent microscopy. c) MALDI-ToF mass spectrum of rPR3, the measured mass and the sequence range is written above the peaks and the colours match the colours of amino acids in the rPR3 sequence showed in (d).

**Fig S2 - Dose-response curves of inhibitors BAY-85-8501, sivelestat, and α1antitrypsin measured with hNE, hPR3, and rPR3.** The activity measurements for each compound concentration were conducted in duplicates or triplicates. For a given inhibitor, each data point on the curve represents the average of two or three replicates and the error bars represent the standard deviation. The relative activity is the ratio of enzymatic activity in the presence of inhibitor to the activity of enzyme in the absence of the inhibitor. The non-linear fit was performed using GraphPad Prism® with a non-linear fit with a standard slope (Hill slope = −1).

**Fig S3 - Immobilization of biotinylated rPR3 (a) on the flow cell 2 and (b) the reference protein δMtDXR on the flow cell 1 of a NAHLC SPR chip.** Injection of protein was followed by washing the unbound protein with the running buffer. In the reference flow cell, a second injection was carried out for a shorter period to increase the protein binding level on the flow cell 1.

**Fig S4 - Analysis of sensorgrams to determine K_D_ values.** Steady state analysis of sensorgrams for BAY-85-8501 (a, K_D_ = 0.03 ± 0.001 μM) and alvelestat (b, K_D_ = 13.5 ± 5.6 μM) binding to rPR3. c) Fitting of BAY85-8501 sensorgrams to the 1:1 ligand binding model for the kinetics analysis with Biacore evaluation software. All graphs were created from data from a single experiment using the Biacore T200 Evaluation Software 3.0.

