## Supplementary information for "Expression and purification of human neutrophil proteinase 3 from insect cells and characterization of ligand binding"

#### Vector constructs

The sequence of PR3 was retrieved from the NCBI in the FASTA format:

```
>NP_002768.3 myeloblastin precursor [Homo sapiens]
MAHRPPSPALASVLLALLLSGAARAAEIVGGHEAQPHSRPYMASLQMRGNPGSHFCGGTLIHPSFVLTAA
HCLRDIPQRLVNVVLGAHNVRTQEPTQQHFSVAQVFLNNYDAENKLNDVLLIQLSSPANLSASVATVQLP
QQDQPVPHGTQCLAMGWGRVGAHDPPAQVLQELNVTVVTFFCRPHNICTFVPRRKAGICFGDSGGPLICD
GIIQGIDSFVIWGCATRLFPDFFTRVALYVDWIRSTLRRVEAKGRP
```

#### The protein sequence and the gene inserts that were embedded in the cloning site EcoRI/HindIII of pFastBac1:

### P2:

Protein sequence:

MKFLVNVALVFMVVYISYIYAMAHRPPSPALASVLLALLLSGAARAAEIVGGHEAQPHSRPYMASLQMRGNPGSHF
CGGTLIHPSFVLTAAHCLRDIPQRLVNVVLGAHNVRTQEPTQQHFSVAQVFLNNYDAENKLNDVLLIQLSSPANLSA
SVATVQLPQQDQPVPHGTQCLAMGWGRVGAHDPPAQVLQELNVTVVTFFCRPHNICTFVPRRKAGICFGDSGGP
LICDGIIQGIDSFVIWGCATRLFPDFFTRVALYVDWIRSTLRRVEAKGRPENLYFQGSAWSHPQFEKGGGSGGGSGG
SAWSHPQFEK

Gene sequence:

GAATTCATGAAGTTCTTGGTCAACGTCGCCTTGGTTTTCATGGTCGTCTACATCAGCTACATTTACGCTATGGCAC
ACCGTCCTCCGTCACCGGCTCTGGCCTCCGTGCTGCTGGCTCTGCTCCTGTCTGGAGCTGCTCGTGCTGCTGAGA
TCGTCGGTGGCCACGAAGCTCAGCCCCACTCTCGCCCATACATGGCCTCACTGCAGATGCGTGGAAACCCAGGT
TCTCACTTCTGCGGAGGTACCCTGATCCACCCTTCATTCGTGCTGACTGCTGCCCACTGCCTGCGCGACATCCCAC
AGCGTCTGGTGAACGTGGTCCTGGGTGCTCACAACGTCAGGACCCAGGAGCCTACTCAGCAGCACTTCTCTGTG
GCTCAGGTCTTCCTGAACAACTACGACGCCGAAAACAAGCTGAACGACGTCCTGCTGATCCAGCTGTCCAGCCC
CGCTAACCTGTCTGCTTCAGTGGCCACCGTCCAGCTGCCACAGCAGGACCAGCCAGTGCCTCACGGCACACAAT
GCCTGGCTATGGGTTGGGGAAGGGTGGGAGCTCACGACCCTCCCGCTCAGGTGCTGCAGGAGCTGAACGTCA
CCGTGGTCACTTTCTTCTGCCGTCCTCACAACATCTGCACCTTCGTGCCCCGCCGTAAGGCTGGCATCTGCTTCG
GAGACTCCGGCGGACCCCTGATCTGCGACGGTATCATCCAGGGCATCGACAGCTTCGTCATCTGGGGTTGCGCT
ACCAGGCTGTTCCCTGACTTCTTCACTAGAGTGGCCCTGTACGTCGACTGGATCAGGTCCACTCTGAGGAGAGT
GGAGGCTAAGGGAAGACCTGAAAACCTGTACTTCCAGGGTTCCGCCTGGAGCCACCCCCAGTTCGAAAAGGG
CGGCGGTAGCGGCGGCGGCAGCGGCGGCTCAGCGTGGTCGCATCCCCAGTTCGAGAAGTAAAAGCTT

### P3:

Protein sequence:

MKFLVNVALVFMVVYISYIYAMAHRPPSPALASVLLALLLSGAARAAEIVGGHEAQPHSRPYMASLQMRGNPGSHF
CGGTLIHPSFVLTAAHCLRDIPQRLVNVVLGAHNVRTQEPTQQHFSVAQVFLNNYDAENKLNDVLLIQLSSPANLSA
SVATVQLPQQDQPVPHGTQCLAMGWGRVGAHDPPAQVLQELNVTVVTFFCRPHNICTFVPRRKAGICFGDSGGP
LICDGIIQGIDSFVIWGCATRLFPDFFTRVALYVDWIRSTLRENLYFQGSAWSHPQFEKGGGSGGGSGGSAWSHPQF
EK

Gene sequence:

GAATTCATGAAGTTCTTGGTCAACGTCGCCTTGGTTTTCATGGTCGTCTACATCAGCTACATTTACGCTATGGCAC
ACCGTCCTCCGTCACCGGCTCTGGCCTCCGTGCTGCTGGCTCTGCTCCTGTCTGGAGCTGCTCGTGCTGCTGAGA
TCGTCGGTGGCCACGAAGCTCAGCCCCACTCTCGCCCATACATGGCCTCACTGCAGATGCGTGGAAACCCAGGT
TCTCACTTCTGCGGAGGTACCCTGATCCACCCTTCATTCGTGCTGACTGCTGCCCACTGCCTGCGCGACATCCCAC
AGCGTCTGGTGAACGTGGTCCTGGGTGCTCACAACGTCAGGACCCAGGAGCCTACTCAGCAGCACTTCTCTGTG
GCTCAGGTCTTCCTGAACAACTACGACGCCGAAAACAAGCTGAACGACGTCCTGCTGATCCAGCTGTCCAGCCC

CGCTAACCTGTCTGCTTCAGTGGCCACCGTCCAGCTGCCACAGCAGGACCAGCCAGTGCCTCACGGCACACAAT
GCCTGGCTATGGGTTGGGGAAGGGTGGGAGCTCACGACCCTCCCGCTCAGGTGCTGCAGGAGCTGAACGTCA
CCGTGGTCACTTTCTTCTGCCGTCCTCACAACATCTGCACCTTCGTGCCCCGCCGTAAGGCTGGCATCTGCTTCG
GAGACTCCGGCGGACCCCTGATCTGCGACGGTATCATCCAGGGCATCGACAGCTTCGTCATCTGGGGTTGCGCT
ACCAGGCTGTTCCCTGACTTCTTCACTAGAGTGGCCCTGTACGTCGACTGGATCAGGTCCACTCTGAGGGAAAA
CCTGTACTTCCAGGGTTCCGCCTGGAGCCACCCCCAGTTCGAAAAGGGCGGCGGTAGCGGCGGCGGCAGCGG
CGGCTCAGCGTGGTCGCATCCCCAGTTCGAGAAGTAAAAGCTT

## P4:

Protein sequence:

MKFLVNVALVFMVVYISYIYAMAHRPPSPALASVLLALLLSGAARAAEIVGGHEAQPHSRPYMASLQMRGNPGSHF
CGGTLIHPSFVLTAAHCLRDIPQRLVNVVLGAHNVRTQEPTQQHFSVAQVFLNNYDAENKLNDVLLIQLSSPANLSA
SVATVQLPQQDQPVPHGTQCLAMGWGRVGAHDPPAQVLQELNVTVVTFFCRPHNICTFVPRRKAGICFGDSGGP
LICDGIIQGIDSFVIWGCATRLFPDFFTRVALYVDWIRSTLRRVEAKGRPENLYFQGHHHHHH

Gene sequence:

GAATTCATGAAGTTCTTGGTCAACGTCGCCTTGGTTTTCATGGTCGTCTACATCAGCTACATTTACGCTATGGCAC
ACCGTCCTCCGTCACCGGCTCTGGCCTCCGTGCTGCTGGCTCTGCTCCTGTCTGGAGCTGCTCGTGCTGCTGAGA
TCGTCGGTGGCCACGAAGCTCAGCCCCACTCTCGCCCATACATGGCCTCACTGCAGATGCGTGGAAACCCAGGT
TCTCACTTCTGCGGAGGTACCCTGATCCACCCTTCATTCGTGCTGACTGCTGCCCACTGCCTGCGCGACATCCCAC
AGCGTCTGGTGAACGTGGTCCTGGGTGCTCACAACGTCAGGACCCAGGAGCCTACTCAGCAGCACTTCTCTGTG
GCTCAGGTCTTCCTGAACAACTACGACGCCGAAAACAAGCTGAACGACGTCCTGCTGATCCAGCTGTCCAGCCC
CGCTAACCTGTCTGCTTCAGTGGCCACCGTCCAGCTGCCACAGCAGGACCAGCCAGTGCCTCACGGCACACAAT
GCCTGGCTATGGGTTGGGGAAGGGTGGGAGCTCACGACCCTCCCGCTCAGGTGCTGCAGGAGCTGAACGTCA
CCGTGGTCACTTTCTTCTGCCGTCCTCACAACATCTGCACCTTCGTGCCCCGCCGTAAGGCTGGCATCTGCTTCG
GAGACTCCGGCGGACCCCTGATCTGCGACGGTATCATCCAGGGCATCGACAGCTTCGTCATCTGGGGTTGCGCT
ACCAGGCTGTTCCCTGACTTCTTCACTAGAGTGGCCCTGTACGTCGACTGGATCAGGTCCACTCTGAGGAGAGT
GGAGGCTAAGGGAAGACCTGAAAACCTGTACTTCCAGGGTCATCATCACCATCACCATTAAAAGCTT

## P5:

Protein sequence:

MKFLVNVALVFMVVYISYIYAHHHHHHENLYFQGIVGGHEAQPHSRPYMASLQMRGNPGSHFCGGTLIHPSFVLTA
AHCLRDIPQRLVNVVLGAHNVRTQEPTQQHFSVAQVFLNNYDAENKLNDVLLIQLSSPANLSASVATVQLPQQDQP
VPHGTQCLAMGWGRVGAHDPPAQVLQELNVTVVTFFCRPHNICTFVPRRKAGICFGDSGGPLICDGIIQGIDSFVI
WGCATRLFPDFFTRVALYVDWIRSTLR

Gene sequence:

GAATTCATGAAGTTCTTGGTCAACGTCGCCTTGGTTTTCATGGTCGTCTACATCAGCTACATTTACGCTCATCATCA
CCATCACCATGAAAACCTGTACTTCCAGGGTATCGTCGGTGGCCACGAAGCTCAGCCCCACTCTCGCCCATACAT
GGCCTCACTGCAGATGCGTGGAAACCCAGGTTCTCACTTCTGCGGAGGTACCCTGATCCACCCTTCATTCGTGCT
GACTGCTGCCCACTGCCTGCGCGACATCCCACAGCGTCTGGTGAACGTGGTCCTGGGTGCTCACAACGTCAGGA
CCCAGGAGCCTACTCAGCAGCACTTCTCTGTGGCTCAGGTCTTCCTGAACAACTACGACGCCGAAAACAAGCTG
AACGACGTCCTGCTGATCCAGCTGTCCAGCCCCGCTAACCTGTCTGCTTCAGTGGCCACCGTCCAGCTGCCACAG
CAGGACCAGCCAGTGCCTCACGGCACACAATGCCTGGCTATGGGTTGGGGAAGGGTGGGAGCTCACGACCCTC
CCGCTCAGGTGCTGCAGGAGCTGAACGTCACCGTGGTCACTTTCTTCTGCCGTCCTCACAACATCTGCACCTTCG
TGCCCCGCCGTAAGGCTGGCATCTGCTTCGGAGACTCCGGCGGACCCCTGATCTGCGACGGTATCATCCAGGGC
ATCGACAGCTTCGTCATCTGGGGTTGCGCTACCAGGCTGTTCCCTGACTTCTTCACTAGAGTGGCCCTGTACGTC
GACTGGATCAGGTCCACTCTGAGGTAAAAGCTT

## P6:

Protein sequence:

MKFLVNVALVFMVVYISYIYAIVGGHEAQPHSRPYMASLQMRGNPGSHFCGGTLIHPSFVLTAAHCLRDIPQRLVNV
VLGAHNVRTQEPTQQHFSVAQVFLNNYDAENKLNDVLLIQLSSPANLSASVATVQLPQQDQPVPHGTQCLAMGW
GRVGAHDPPAQVLQELNVTVVTFFCRPHNICTFVPRRKAGICFGDSGGPLICDGIIQGIDSFVIWGCATRLFPDFFTR
VALYVDWIRSTLRENLYFQGSAWSHPQFEKGGGSGGGSGGSAWSHPQFEK

Gene sequence:

GAATTCATGAAGTTCTTGGTCAACGTCGCCTTGGTTTTCATGGTCGTCTACATCAGCTACATTTACGCTATCGTCG
GTGGCCACGAAGCTCAGCCCCACTCTCGCCCATACATGGCCTCACTGCAGATGCGTGGAAACCCAGGTTCTCAC
TTCTGCGGAGGTACCCTGATCCACCCTTCATTCGTGCTGACTGCTGCCCACTGCCTGCGCGACATCCCACAGCGT
CTGGTGAACGTGGTCCTGGGTGCTCACAACGTCAGGACCCAGGAGCCTACTCAGCAGCACTTCTCTGTGGCTCA
GGTCTTCCTGAACAACTACGACGCCGAAAACAAGCTGAACGACGTCCTGCTGATCCAGCTGTCCAGCCCCGCTA
ACCTGTCTGCTTCAGTGGCCACCGTCCAGCTGCCACAGCAGGACCAGCCAGTGCCTCACGGCACACAATGCCTG
GCTATGGGTTGGGGAAGGGTGGGAGCTCACGACCCTCCCGCTCAGGTGCTGCAGGAGCTGAACGTCACCGTG
GTCACTTTCTTCTGCCGTCCTCACAACATCTGCACCTTCGTGCCCCGCCGTAAGGCTGGCATCTGCTTCGGAGAC
TCCGGCGGACCCCTGATCTGCGACGGTATCATCCAGGGCATCGACAGCTTCGTCATCTGGGGTTGCGCTACCAG
GCTGTTCCCTGACTTCTTCACTAGAGTGGCCCTGTACGTCGACTGGATCAGGTCCACTCTGAGGGAAAACCTGTA
CTTCCAGGGTTCCGCCTGGAGCCACCCCCAGTTCGAAAAGGGCGGCGGTAGCGGCGGCGGCAGCGGCGGCTC
AGCGTGGTCGCATCCCCAGTTCGAGAAGTAAAAGCTT

### PCR primers

The M13 primers for the PCR reactions with the bacmid sequence were obtained from Genscript. (Forward primer: 5′-CCCAGTCACGACGTTGTAAAACG-3′, reverse primer: 5′-AGCGGATAACAATTTCACACAGG-3′).

### Supplementary figures

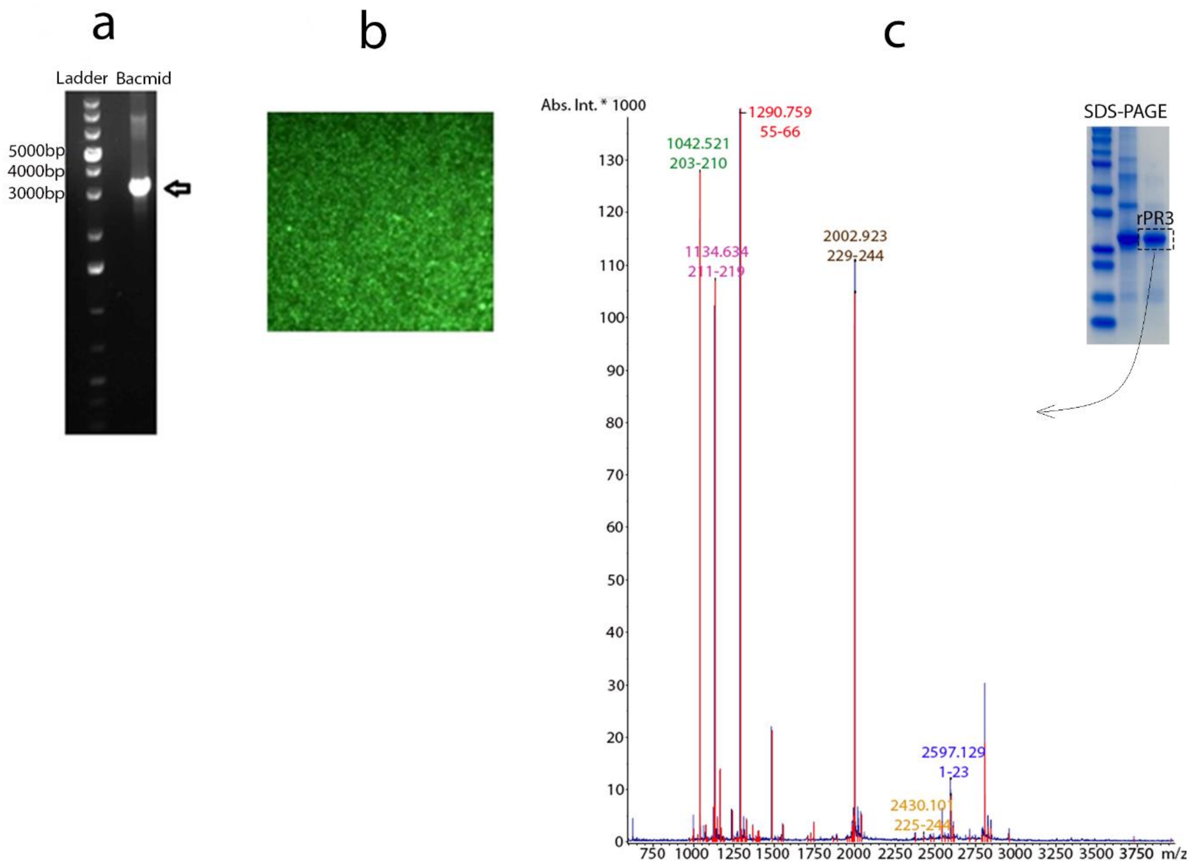


d

```
AEIVGGHEAQPHSRPYMASLQMRGNPGSHFCGGTLIHPSFVLTAAHCLRDIPQRLVNVVLGAHNVRTQEPTQQ
HFSVAQVFLNNYDAENKLNDVLLIQLSSPANLSASVATVQLPQQDQPVPHGTQCLAMGWGRVGAHDPPAQVLQ
ELNVTVVTFFCRPHNICTFVPRRKAGICFGDSGGPLICDGIIQGIDSFVIWGCATRLFPDFFTRVALYVDWIR
STLRRVEAKGRPENLYFQGHHHHHH
```

**Fig S1- Expression and identification of rPR3.** a) Agarose gel to detect bacmid production. b) Sf9 cells infected with baculovirus showing expression of YFP as green dots using fluorescent microscopy. c) MALDI-ToF mass spectrum of rPR3. The rPR3 band was cut from the SDS-PAGE, digested with trypsin and used for the MS study. The peaks are labelled with the measured mass and the sequence range whereas the font colour corresponds to the coloured amino acids in the rPR3 sequence shown in (d).

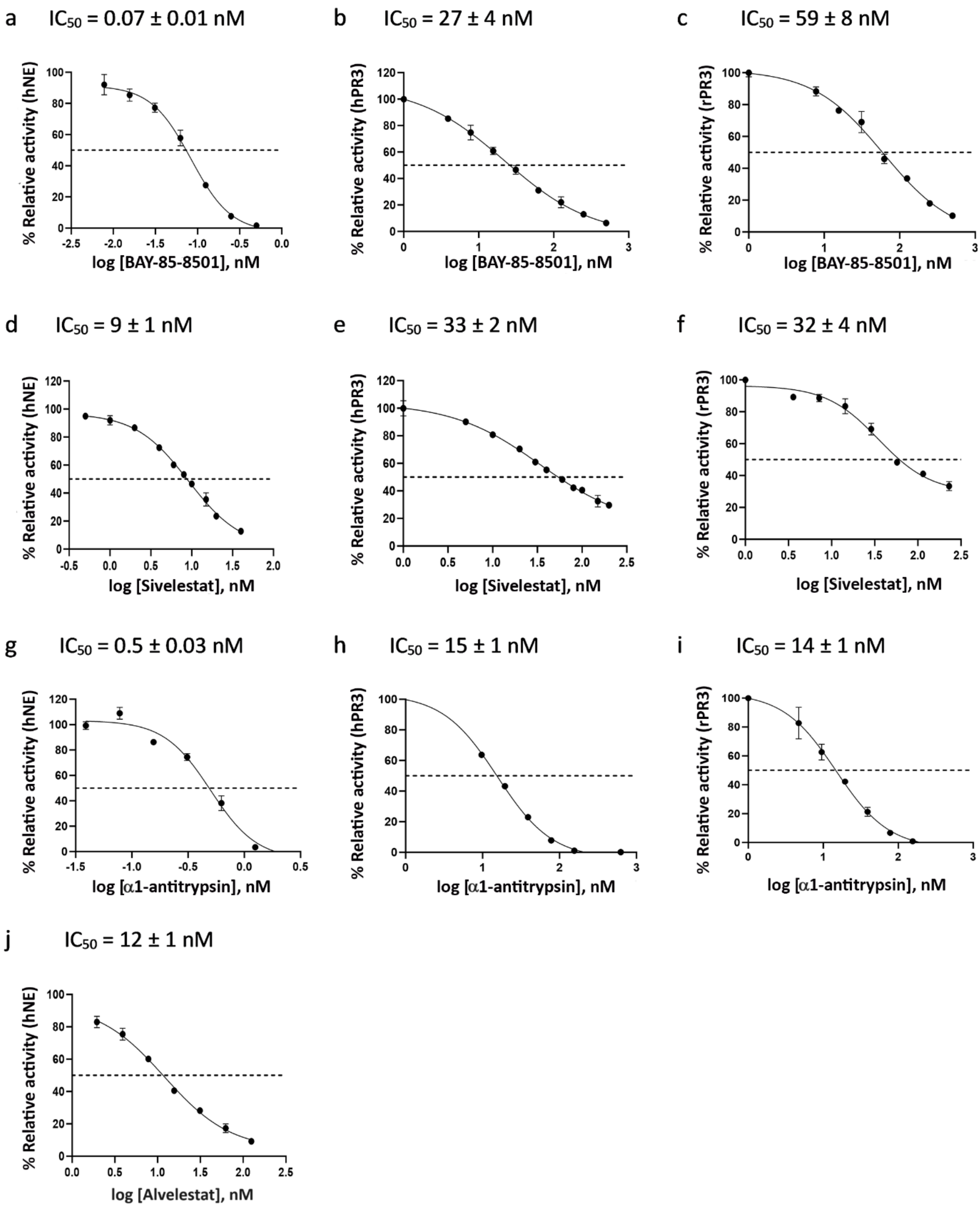


**Fig S2- Dose-response curves of inhibitors BAY-85-8501, sivelestat, and α1antitrypsin measured with hNE, hPR3, and rPR3.** The activity measurements for each compound concentration were conducted in duplicates or triplicates. For a given inhibitor, each data point on the curve represents the average of two or three replicates and the error bars represent the standard deviation. The relative activity is the ratio of enzymatic activity in the presence of inhibitor to the activity of enzyme in the absence of the inhibitor. The non-linear fit was performed using GraphPad Prism® with a non-linear fit with a standard slope (Hill slope = -1).

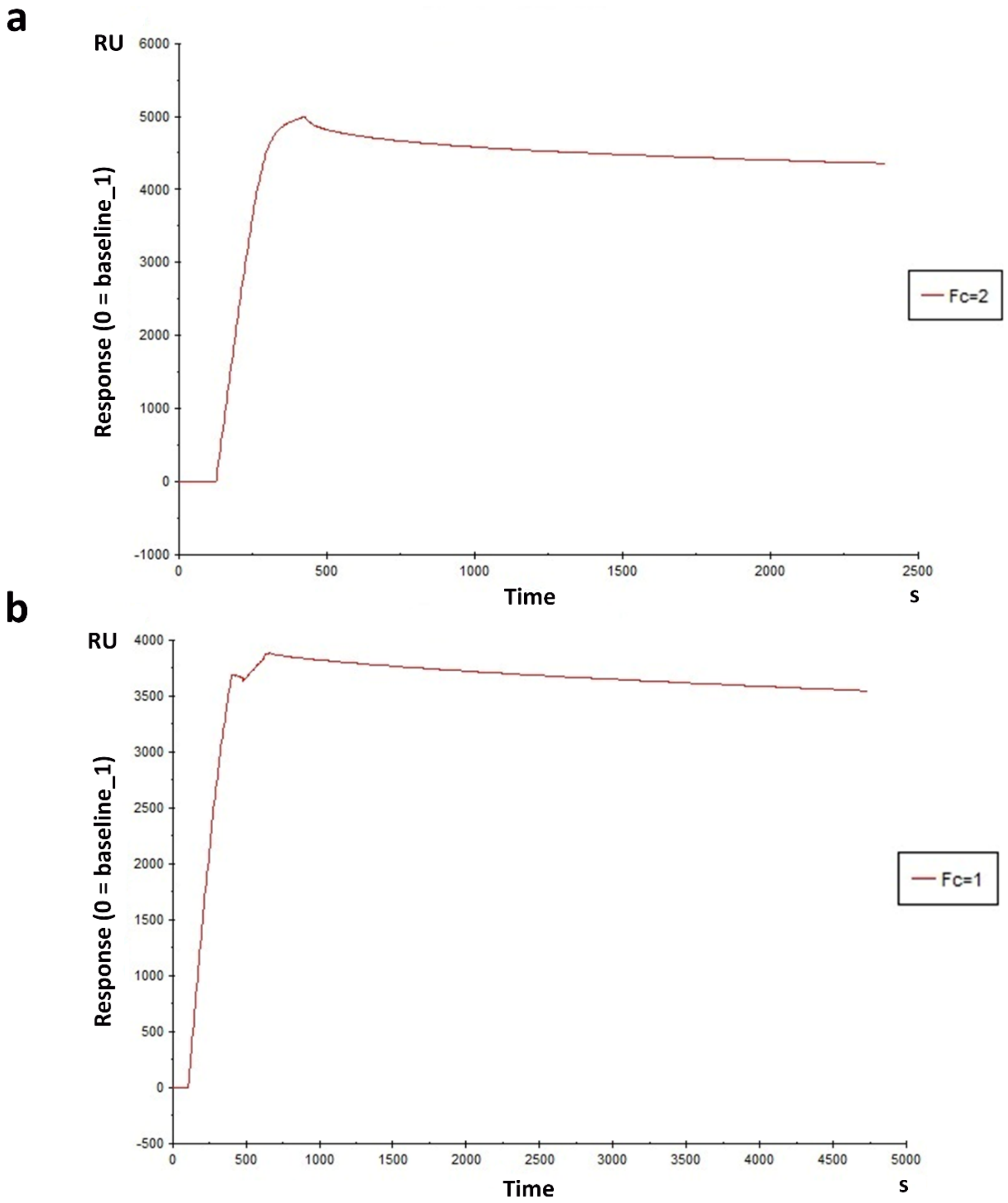


**Fig S3- Immobilization of biotinylated rPR3 (a) on the flow cell 2 and (b) the reference protein δMtDXR on the flow cell 1 of a NAHLC SPR chip.** Injection of protein was followed by washing the unbound protein with the running buffer. In the reference flow cell, a second injection was carried out for a shorter period to increase the protein binding level on the flow cell 1.

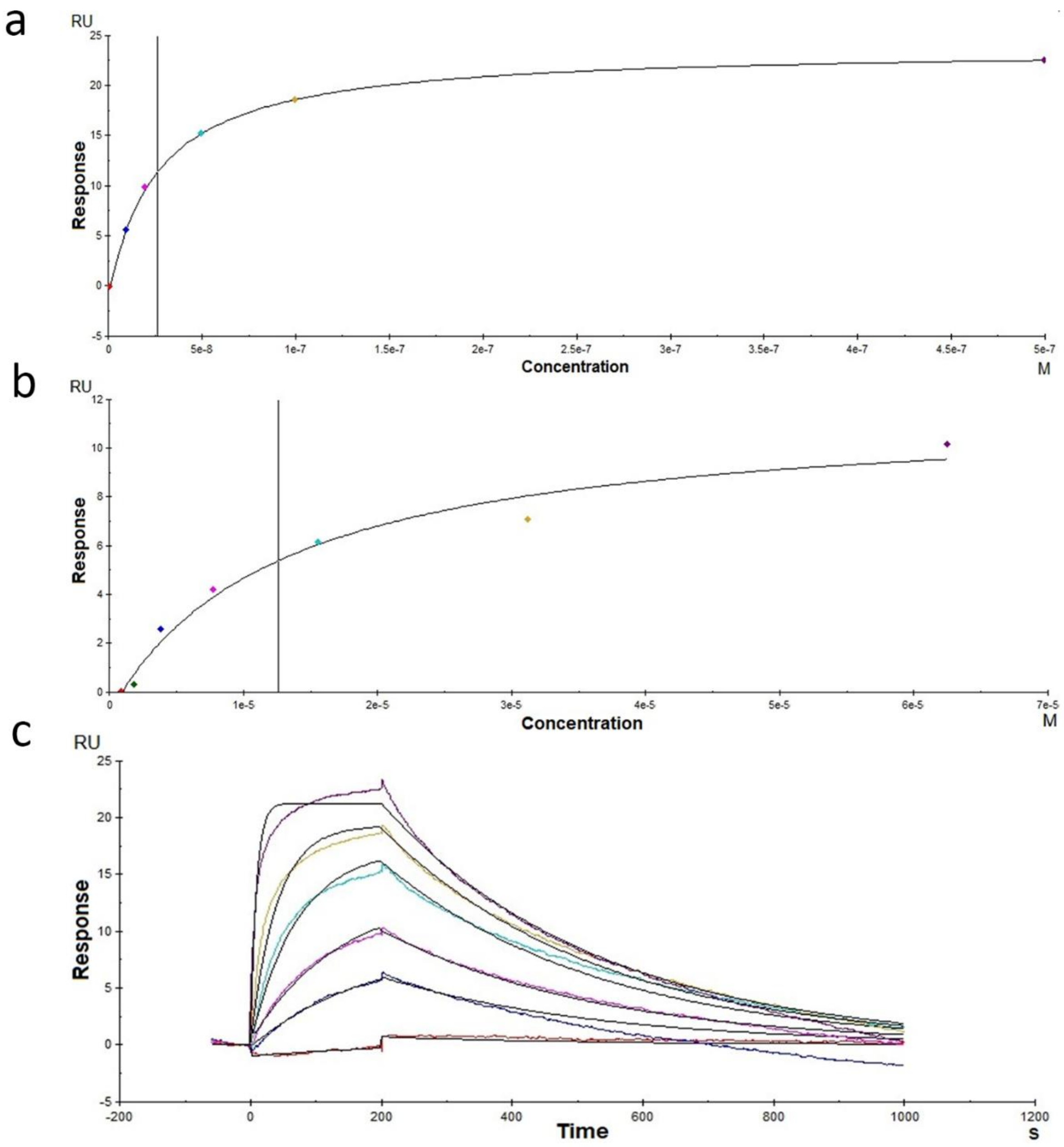


**Fig S4- Analysis of sensorgrams to determine $K_D$ Values.** Steady state analysis of sensorgrams for BAY-85-8501 (a, $K_D$ = 0.03 ± 0.001 $\mu$M) and alvelestat (b, $K_D$ = 13.5 ± 5.6 $\mu$M) binding to rPR3. c) Fitting of BAY85-8501 sensorgrams to the 1:1 ligand binding model for the kinetics analysis with Biacore evaluation software. All graphs were created from data from a single experiment using the Biacore T200 Evaluation Software 3.0.
